# *SSD1* suppresses phenotypes induced by the lack of Elongator-dependent tRNA modifications

**DOI:** 10.1101/596197

**Authors:** Fu Xu, Anders S. Byström, Marcus J.O. Johansson

## Abstract

The Elongator complex promotes formation of 5-methoxycarbonylmethyl (mcm^5^) and 5-carbamoylmethyl (ncm^5^) side-chains on uridines at the wobble position of cytosolic eukaryotic tRNAs. In all eukaryotic organisms tested to date, the inactivation of Elongator not only leads to the lack of mcm^5^/ncm^5^ groups in tRNAs, but also a wide variety of phenotypes. Although the phenotypes are most likely caused by a translational defect induced by reduced functionality of the hypomodified tRNAs, the mechanism(s) underlying individual phenotypes are poorly understood. In this study, we show that the genetic background modulates the phenotypes induced by the lack of mcm^5^/ncm^5^ groups in *Saccharomyces cerevisiae*. We show that the stress-induced growth defects of Elongator mutants are stronger in the W303 than in the closely related S288C genetic background and that the phenotypic differences are caused by the known polymorphism at the locus for the mRNA binding protein Ssd1. Moreover, the mutant *ssd1* allele found in W303 cells is required for the reported histone H3 acetylation and telomeric gene silencing defects of Elongator mutants. The difference at the *SSD1* locus also partially explains why the simultaneous lack of mcm^5^ and 2-thio groups at wobble uridines is lethal in the W303 but not in the S288C background. Collectively, our results demonstrate that the *SSD1* locus modulates phenotypes induced by the lack of Elongator-dependent tRNA modifications.

**Author Summary:** Modified nucleosides in the anticodon region of tRNAs are important for the efficiency and fidelity of translation. The Elongator complex promotes formation of several related modified uridine residues at the wobble position of eukaryotic tRNAs. In yeast, plants, worms, mice and humans, mutations in genes for Elongator subunits lead to a wide variety of different phenotypes. Here, we show that the genetic background modulates the phenotypic consequences of the inactivation of budding yeast Elongator. This background effect is largely a consequence of a polymorphism at the *SSD1* locus, encoding a RNA binding protein that influences translation, stability and/or localization of mRNAs. We show that several phenotypes reported for yeast Elongator mutants are either significantly stronger or only detectable in strains harboring a mutant *ssd1* allele. Thus, *SSD1* is a suppressor of the phenotypes induced by the hypomodification of tRNAs.

## Introduction

A general feature of tRNA molecules is that a subset of their nucleosides harbors post-transcriptional modifications. Modified nucleosides are frequently found in the anticodon region of tRNAs, especially at position 34 (the wobble nucleoside) and 37. Modifications at these positions typically influence the decoding properties of tRNAs by improving or restricting anticodon-codon interactions [1, 2]. Uridines present at the wobble position in eukaryotic cytoplasmic tRNAs often harbor a 5-methoxycarbonylmethyl (mcm^5^) or 5-carbamoylmethyl (ncm^5^) side-chain and sometimes also a 2-thio (s^2^) or 2’-*O*-methyl group [3, 4]. The first step in the synthesis of the mcm^5^ and ncm^5^ side-chains requires the Elongator complex, which is composed of six Elp proteins (Elp1-Elp6) [5–9]. Elongator is thought to catalyze the addition of a carboxymethyl (cm) group to the 5-position of the uridine which is then converted to mcm^5^ by the Trm9/Trm112 complex or to ncm^5^ by a yet unidentified mechanism [5, 8–12].

In the budding yeast *Saccharomyces cerevisiae*, the inactivation of any of the six *ELP* genes (*ELP1-ELP6*) not only leads to the lack of the mcm^5^/ncm^5^ groups but also slower growth rate and numerous additional phenotypes [5, 13]. These phenotypes include increased sensitivity to elevated temperatures and various chemical stresses as well as defects in transcription, exocytosis, telomeric gene silencing, and protein homeostasis [14–18]. Even though Elongator mutants lack mcm^5^/ncm^5^ groups in 11 tRNA species [5, 19], the pleiotropic phenotypes are suppressed by increased expression of various combinations of the hypomodified forms of the three tRNA species that normally harbor a mcm^5^s^2^U_34_ residue, 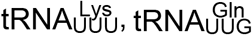 and 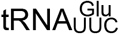 [18, 20, 21]. These findings suggest the pleiotropic phenotypes of Elongator mutants are caused by a reduced functionality of the hypomodified 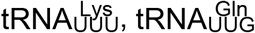 and 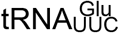 in translation [20, 21]. The importance of the modified wobble residue in these tRNAs was supported by the finding that strains lacking the s^2^ group show the same but slightly weaker phenotypes that are also suppressed by increased expression of the three tRNAs [20, 21]. Moreover, ribosome profiling experiments have shown that the inactivation of Elongator causes an accumulation of ribosomes with AAA, CAA or GAA codons in the ribosomal A-site [18, 22, 23]. However, the pausing at the codons appears to be relatively small [18, 22] and the mechanism(s) underlying the pleiotropic phenotypes of Elongator mutants are poorly understood.

In yeast, the cell wall stress that arises during normal growth or through environmental challenges is sensed and responded to by the cell wall integrity (CWI) pathway [24, 25]. The CWI pathway is induced by several different types of stresses, including growth at elevated temperatures, hypo-osmotic shock, and exposure to various cell wall stressing agents [25]. A family of cell surface sensors (Wsc1-Wsc3, Mid2 and Mtl1) detects the cell wall stress and recruits the guanine nucleotide exchange factors Rom1 and Rom2 which activate the small GTPase Rho1. Rho1-GTP binds and activates several effectors, including the kinase Pkc1. Pkc1 activates a downstream MAP kinase cascade comprised of the MAPKKK Bck1, the two redundant MAPKK Mkk1 and Mkk2, and the MAPK Mpk1 (Slt2). The phosphorylated Mpk1 then activates factors that promote transcription of genes important for cell wall biosynthesis and remodeling.

In addition to the CWI pathway, several other factors and pathways are known to influence the cell wall remodeling that occurs upon stress, e.g. the mRNA-binding protein Ssd1. Ssd1 has been reported to bind and influence the translation, stability and/or localization of a subset of cellular mRNAs of which many encode proteins important for cell wall biosynthesis and remodeling, [26–31]. The wild-type *SSD1* gene was originally identified as a suppressor of the lethality induced by a deletion of the *SIT4* gene, which encodes a phosphatase involved in a wide range of cellular processes [32]. The study also led to the finding that some wild-type *S. cerevisiae* laboratory strains harbor a mutation at the *SSD1* locus that is synthetic lethal with the *sit4Δ* allele [32]. The *SSD1* locus has since been genetically implicated in many cellular processes, including cell wall integrity, various signal transduction pathways, cell morphogenesis, cellular aging, virulence, and transcription by RNA polymerase I, II and III [33–38]. Although the mechanisms by which Ssd1 influences these processes are poorly understood, they possibly involve both direct and indirect effects of Ssd1’s influence on messenger ribonucleoprotein (mRNP) complexes [28, 29, 39]. With respect to the transcripts that encode cell wall remodeling factors, Ssd1 seems to act as a translational repressor and this function is controlled by the protein kinase Cbk1, which is a component in the RAM (Regulation of Ace2 and cellular morphogenesis) network [28]. In addition to relieving the translational repression, the phosphorylation of Ssd1 appears to promote polarized localization of some Ssd1-associated mRNAs [28, 31].

In this study, we show that increased activation of the CWI signaling pathway counteracts the temperature sensitive (Ts) growth defect of *elp3Δ* mutants in the W303 but not in the S288C genetic background. Further, the stress-induced growth phenotypes caused by the tRNA modification defect are generally stronger in W303-than in S288C-derived strains. We show that the phenotypic differences are due to the allelic variation at the *SSD1* locus, i.e. the phenotypes are aggravated by the nonsense *ssd1-d2* allele found in the W303 background. We also show that the phenotypes linking the tRNA modification defect to histone acetylation and telomeric gene silencing are caused by a synergistic interaction with the *ssd1-d2* allele. The difference at the *SSD1* locus also provide a partial explanation to the finding [18, 40, 41] that cells lacking both the mcm^5^ and s^2^ group are viable in the S288C but not in the W303 background.

## Results

### The Ts phenotype of W303-derived *elp3Δ* cells is partially suppressed by increased expression of factors in the CWI signaling pathway

The observation that Elongator mutants are sensitive to cell wall stressing agents, e.g. calcofluor white and caffeine, implies a defect in cell wall integrity [14]. This notion is further supported by the finding that the Ts growth defect of Elongator-deficient cells is partially suppressed by osmotic support (1 M sorbitol) [42]. As the caffeine sensitivity and Ts growth defect are suppressed by increased levels of the hypomodified 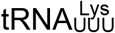 and 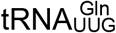 [20], the phenotypes are likely caused by reduced functionality of these tRNAs in translation. To further define the wall integrity defect in Elongator mutants, we determined, in the W303 genetic background, if the Ts phenotype of *elp3Δ* cells is suppressed by increased expression of factors in the CWI signaling pathway. The analyses revealed that the introduction of a high-copy *MID2, WSC2, ROM1*, or *PKC1* plasmid into the *elp3Δ* strain partially suppressed the growth defect at 37°C (Fig 1A). No suppression of the phenotype was observed when the cells carried a high-copy *RHO1, BCK1* or *MPK1* plasmid (Fig 1A). As the overexpression of neither the upstream GTPase (Rho1) nor the downstream kinases (Bck1 and Mpk1) suppressed the Ts phenotype, we considered the possibility that the levels of these factors may be too high when expressed from a high-copy plasmid. Accordingly, low-copy *RHO1, BCK1* and *MPK1* plasmids suppress the Ts phenotype of *elp3Δ* cells to a level similar to that observed for the high-copy *MID2, WSC2, ROM1*, and *PKC1* plasmids (Fig 1A). The level of suppression is, however, smaller than that observed for increased *tK*(*UUU*) and *tQ*(*UUG*) dosage, encoding 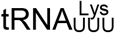 and 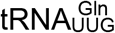, respectively (Fig 1A).

**Fig 1.**
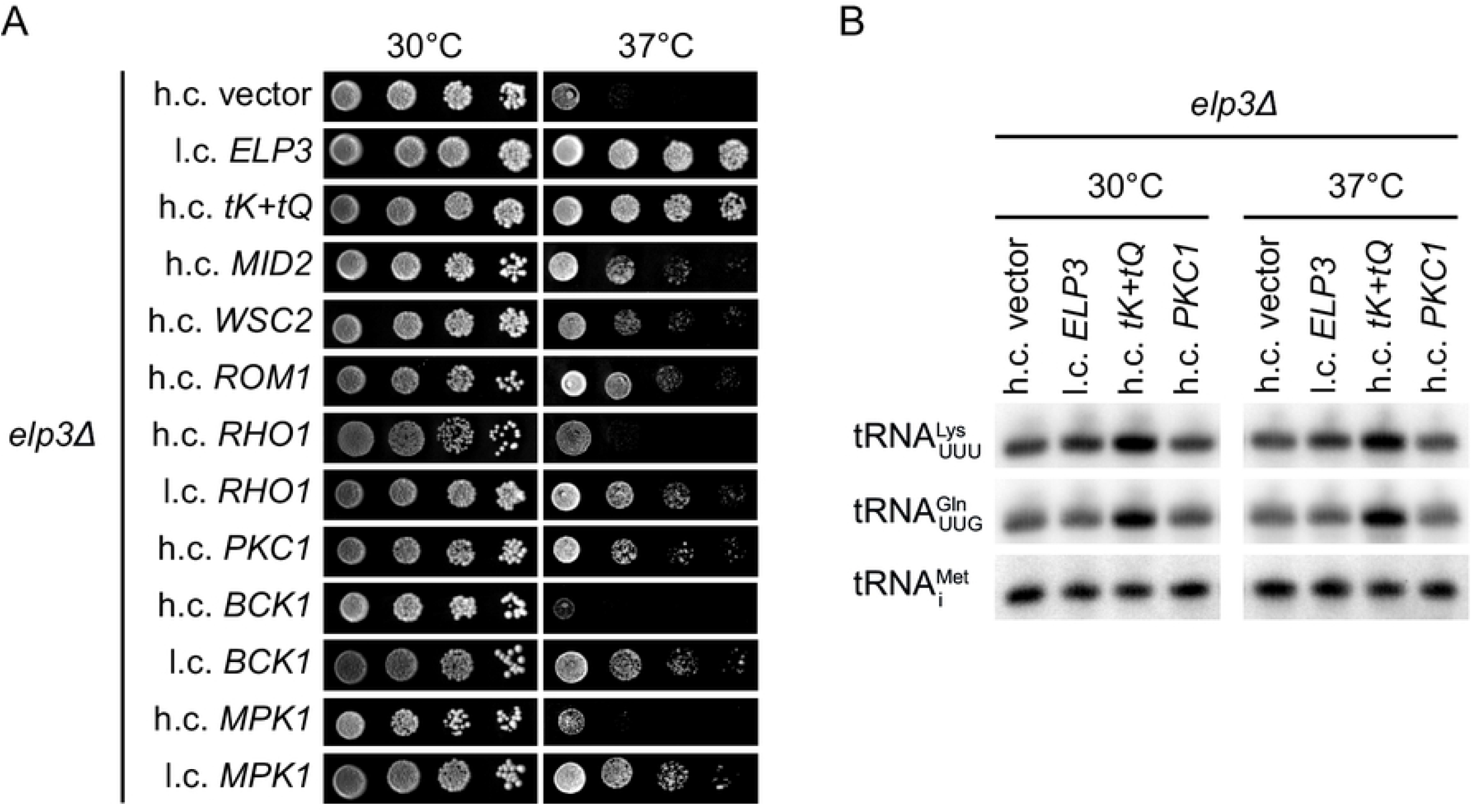
Increased expression of factors in the CWI signaling pathway counteracts the Ts phenotype of W303-derived *elp3Δ* cells. (A) Growth of the *elp3Δ* (UMY3269) strain carrying the indicated high-copy (h.c.) or low-copy (l.c.) *LEU2* plasmids. The high copy plasmid carrying the *tK*(*UUU*) and *tQ*(*UUG*) genes [72] is abbreviated as h.c. *tK+tQ*. Cells were grown over-night at 30°C in liquid synthetic complete medium lacking leucine (SC-leu), serially diluted, spotted on SC-leu plates, and incubated at 30°C or 37°C for 3 days. (B) Northern analysis of total RNA isolated from *elp3Δ* (UMY3269) cells carrying the indicated plasmids. The cells were grown in SC-leu medium at 30°C or 37°C. The blot was probed for 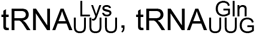 and 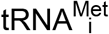 using radiolabeled oligonucleotides. 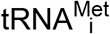 serves as the loading control.

Since the Ts phenotype of *elp3Δ* cells is counteracted by elevated 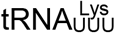 and 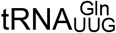 levels [20], it was possible that the activation of the CWI pathway leads to an increase in their relative abundance. To investigate this possibility, we used northern blotting to analyze the effect of increased *PKC1* dosage on the levels of 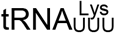 and 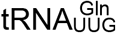 in *elp3Δ* cells grown at either 30°C or 37°C. The blots were also probed for 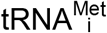, which served as the loading control. In contrast to the ≈2.5-fold increase in 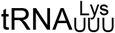 and 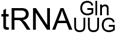 levels induced by increased *tK*(*UUU*) and *tQ*(*UUG*) dosage (Fig 1B and S1 Table), the abundance of the tRNAs were largely unaffected by increased *PKC1* dosage at both 30°C and 37°C (Fig 1B and S1 Table). Collectively, our results suggest that the Ts phenotype of *elp3Δ* cells is, at least in the W303 genetic background, partially suppressed by increased activation of the CWI signaling pathway.

### The allelic variant at the *SSD1* locus modulates the growth phenotypes of *elp3Δ* cells

Phenotypes caused by a mutation in an individual gene can be modulated by the genetic background of the cell. In fact, the Ts phenotype induced by an *elp3Δ* allele is more pronounced in strains derived from W303 than in those from S288C (Fig 2A). As the inactivation of Elongator causes a lack of wobble mcm^5^/ncm^5^ groups in both strain backgrounds [5, 43], the Ts phenotype is likely modulated by genetic variation between W303 and S288C. The difference in phenotype (Fig 2A) prompted us to investigate if increased *MID2, WSC2, ROM1, RHO1, PKC1, BCK1* or *MPK1* dosage counteracts the Ts phenotype of *elp3Δ* cells in the S288C background. The analyses showed that none of the plasmids counteracted the phenotype (S1A Fig). Moreover, the growth defect of *elp3Δ* cells at 37°C is counteracted by osmotic support (1 M sorbitol) in the W303, but not in the S288C background (S1B Fig). Importantly, the Ts phenotype of *elp3Δ* cells is suppressed by increased *tK*(*UUU*) and *tQ*(*UUG*) dosage in both genetic backgrounds (Fig 1A and S1A Fig), showing that the underlying cause is the hypomodified 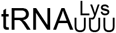 and 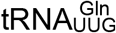. We conclude that the genetic background influences the phenotypes linking the tRNA modification defect to cell wall integrity.

**Fig 2.**
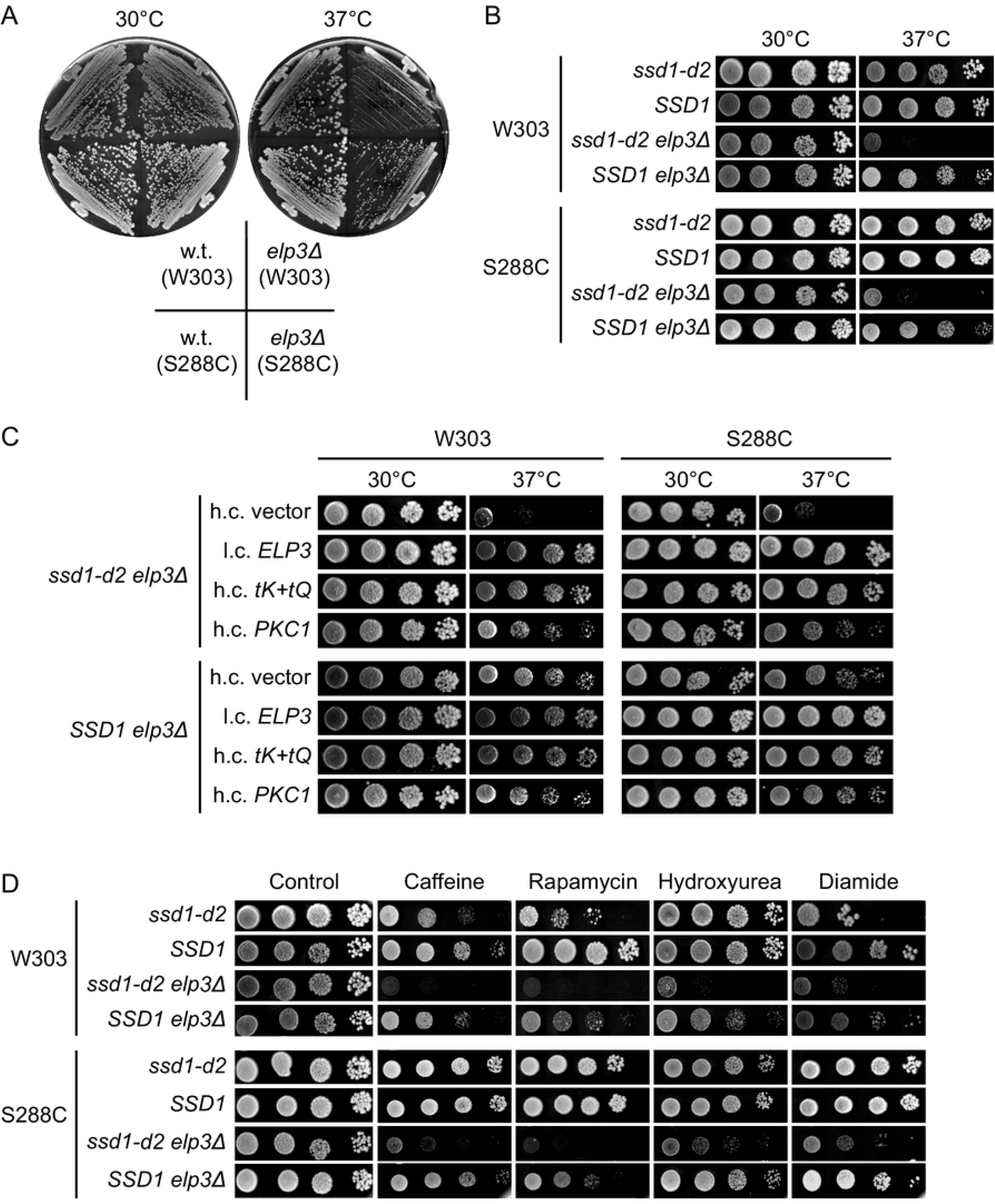
The growth phenotypes *elp3Δ* cells are modulated by the allele at the *SSD1* locus. (A) Growth of *elp3Δ* mutants in the W303 and S288C genetic backgrounds. The wild-type (W303-1A and BY4741) and *elp3Δ* strains (UMY3269 and MJY1036) were streaked on SC plates and incubated at 30°C or 37°C for 3 days. (B) Effects of the *ssd1-d2/SSD1* alleles on the growth of *elp3Δ* strains in the W303 and S288C genetic backgrounds. The *ssd1-d2* (W303-1A and UMY4432), *SSD1* (UMY3385 and BY4741), *ssd1-d2 elp3Δ* (UMY3269 and UMY4439) and *SSD1 elp3Δ* (UMY4456 and MJY1036) strains were grown over-night at 30°C in liquid SC medium, serially diluted, spotted on SC plates, and incubated at 30°C or 37°C for 3 days. (C) Effects of increased *PKC1* dosage on the growth of *elp3Δ ssd1-d2* and *elp3Δ SSD1* strains. The relevant strains (UMY3269,UMY4456, UMY4439, and MJY1036) carrying the indicated plasmids were grown over-night at 30°C in liquid SC-leu medium, serially diluted, spotted on SC-leu plates, and incubated for 3 days at 30°C or 37°C. (D) Influence of the *ssd1-d2* allele on growth phenotypes induced by various stress-inducing agents. The strains from B were grown over-night at 30°C in liquid SC medium, serially diluted, spotted on SC plates and SC plates supplemented with caffeine, rapamycin, hydroxyurea or diamide. The plates were incubated for 3 days at 30°C.

Although W303 is closely related to S288C, comparisons of the genomes identified polymorphisms in ≈800 genes that lead to variations in the amino acid sequence [44, 45]. To identify the cause of the phenotypic differences between the *elp3Δ* strains, we examined polymorphisms that have been shown to be physiologically relevant. The polymorphism at the *SSD1* locus was a good candidate as *SSD1* has been genetically implicated in many cellular processes, including the maintenance of cellular integrity [32, 33, 46–50]. Ssd1 is a RNA binding protein that associates with a subset of mRNAs of which many encode proteins important for cell wall biosynthesis and remodeling [26–28]. The *SSD1* allele in the S288C background encodes the full-length Ssd1 protein (1250 amino acids) whereas the allele in W303, designated *ssd1-d2*, contains a nonsense mutation that introduces a premature stop codon at the 698^th^ codon of the open reading frame [32, 35]. To investigate if the allele at the *SSD1* locus contributes to the phenotypic differences between the *elp3Δ* mutants, we analyzed congenic *ssd1-d2, SSD1, ssd1-d2 elp3Δ*, and *SSD1 elp3Δ* strains in both strain backgrounds. HPLC analyses of the nucleoside composition of total tRNA from these strains showed that the levels of ncm^5^U, mcm^5^U, and mcm^5^s^2^U are comparable in the *ssd1-d2* and *SSD1* strains and not detectable in the *ssd1-d2 elp3Δ* and *SSD1 elp3Δ* mutants (S2 Table). The HPLC analyses also showed that the allele at the *SSD1* locus does not appear to influence the abundance of other modified nucleosides present in tRNAs (S2 Fig). By analyzing the growth of the strains, we found that the *ssd1-d2* allele augments the Ts phenotype of *elp3Δ* cells in both backgrounds (Fig 2B and S3 Table). The *ssd1-d2* allele also appears to cause a slightly reduced growth rate of strains with a wild-type *ELP3* gene at the elevated temperature (S3 Table). Even though the *elp3Δ* mutants grow slower than the *ELP3* strains at 30°C, the effect of the *ssd1-d2* allele is less pronounced at that temperature (S3 Table). In both genetic backgrounds, the increased activation of the CWI signaling pathway, through increased *PKC1* dosage, suppresses the Ts phenotype of the *ssd1-d2 elp3Δ*, but not the *SSD1 elp3Δ* strains (Fig 2C). Although the *elp3Δ* strains show increased sensitivity to caffeine, irrespective of the allele at the *SSD1* locus, the *ssd1-d2 elp3Δ* cells are in both backgrounds more caffeine-sensitive than the *SSD1 elp3Δ* cells (Fig 2D). This observation is consistent with the previous finding that the *ssd1-d2* allele enhances the growth inhibitory effects of caffeine [32].

The inactivation of Elongator not only leads to increased sensitivity to caffeine, but also to other stress-inducing agents [14, 17, 18]. To investigate if the *ssd1-d2* allele influences these phenotypes, we analyzed the growth of the *ssd1-d2, SSD1, ssd1-d2 elp3Δ*, and *SSD1 elp3Δ* strains on medium containing rapamycin, hydroxyurea or diamide. The analyses revealed that the *ssd1-d2* allele, irrespective of background, also increases the sensitivity of *elp3Δ* cells to these compounds (Fig 2D). Northern blot analyses revealed that the relative abundance of 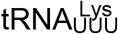 and 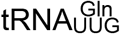 is largely unaffected by allelic variant at the *SSD1* locus (S3 Fig and S4 Table), indicating that the *ssd1-d2* allele influences the phenotypes by a different mechanism. Collectively, these results show that the allele at the *SSD1* locus influences stress-induced growth defects of *elp3Δ* cells.

### The *ssd1-d2* allele is required for the histone acetylation and telomeric gene silencing defects of *elp3Δ* mutants

In the W303 background, *elp3Δ* mutants show reduced acetylation of histone H3 [15]. Although the phenotype was originally thought to reflect a function of Elongator in RNA polymerase II transcription [15], the reduced acetylation of lysine-14 (K14) in histone H3 was subsequently shown to be an indirect consequence of the tRNA modification defect [20]. To investigate if the *ssd1-d2* allele contributes to the phenotype, we analyzed, in the W303 background, the histone H3 K14 acetylation levels in the *ssd1-d2, ssd1-d2 elp3Δ, SSD1*, and *SSD1 elp3Δ* strains. As previously shown [15, 20], the level of K14 acetylation is lower in the *ssd1-d2 elp3Δ* mutant than in the *ssd1-d2* strain (Fig 3A). However, the *SSD1* and *SSD1 elp3Δ* strains show comparable levels of K14 acetylated histone H3, indicating that it is the combination of the *elp3Δ* and *ssd1-d2* alleles that induces the histone H3 acetylation defect.

**Fig 3.**
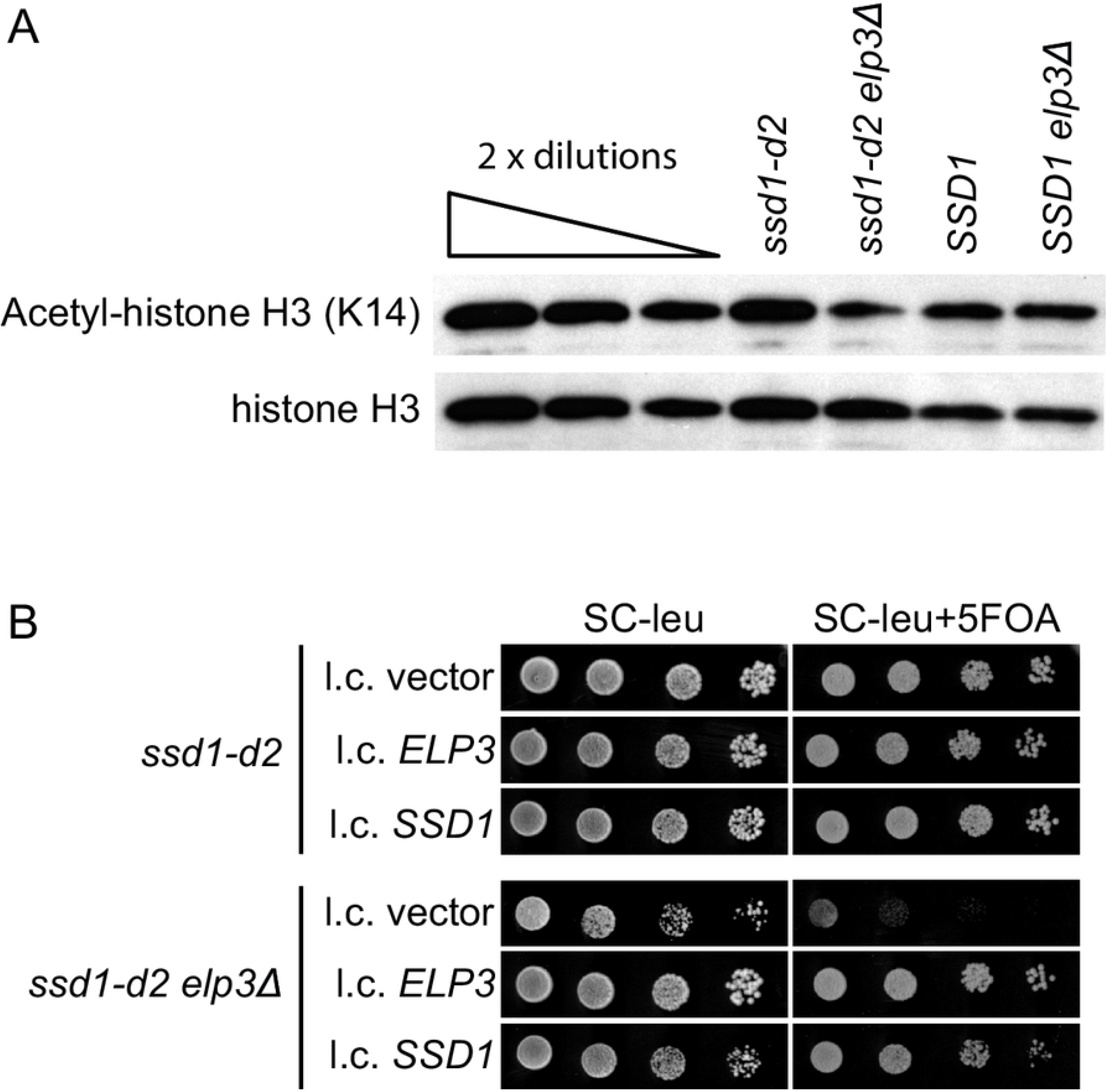
The *ssd1-d2* allele is required for the histone acetylation and telomeric gene silencing defects of *elp3Δ* cells. (A) Western analysis of histones isolated from the *ssd1-d2* (W303-1A), *ssd1-d2 elp3Δ* (UMY3269), *SSD1* (UMY3385) and *SSD1 elp3Δ* (UMY4456) strains grown in SC medium at 30°C. Polyclonal anti-acetyl-histone H3 and anti-histone H3 antibodies were used to detect the indicated proteins. The blot is a representative of two independent experiments. (B) Influence of the *ssd1-d2* allele on telomeric gene silencing in *elp3Δ* cells. The *ssd1-d2 TELVIIL::URA3* (UMY2584) and *ssd1-d2 elp3Δ TELVIIL::URA3* (UMY3790) strains carrying the indicated plasmids were grown over-night at 30°C in liquid SC-leu medium, serially diluted, spotted on SC-leu and SC-leu+5-FOA plates, and incubated for 3 days at 30°C.

W303-derived Elongator mutants have also been reported to show delayed transcriptional activation of the *GAL1* and *GAL10* genes upon a shift from raffinose-to galactose-containing medium [20, 51]. To determine if the *ssd1-d2* allele influences this phenotype, we analyzed the induction of the *GAL1* mRNA in the W303-derived *ssd1-d2, ssd1-d2 elp3Δ, SSD1*, and *SSD1 elp3Δ* strains. Unexpectedly, we observed no obvious delay in the induction of *GAL1* transcripts in the *elp3Δ* strains regardless of the nature of the allele at the *SSD1* locus (S4 Fig). As the phenotype is thought to reflect a reduced ability of Elongator mutants to adapt to new growth conditions [51], it is possible that differences in media or the handling of the cultures can explain why *elp3Δ* cells show rapid *GAL1* induction in our experiments.

Another phenotype reported for Elongator mutants in the W303 background is a defect in telomeric gene silencing [17, 21]. The telomere silencing defect of Elongator mutants was inferred from experiments where the expression of a *URA3* gene integrated close to the left telomere of chromosome VII was assayed [17, 21]. The defect in telomeric gene silencing leads to increased expression of the *URA3* gene, which is scored as reduced growth on plates containing 5-fluoroorotic acid (5-FOA) [17, 21]. Even though the integration of an *SSD1* allele into *ssd1-d2* cells does not influence the level of telomeric gene silencing [36], the inactivation of *SSD1* does increase the expression of a reporter gene at the silent mating type locus HMR [52]. The latter finding implies that the *ssd1-d2* allele may influence the assembly of silent chromatin and consequently contribute to the silencing defect in Elongator mutants. Accordingly, the introduction of a low-copy *SSD1* plasmid into the *ssd1-d2 elp3Δ TELVIIL::URA3* strain [21] complemented the 5-FOA sensitivity to a level similar to that observed with a plasmid carrying the wild-type *ELP3* gene (Fig 3B). Thus, the telomeric gene silencing defect in Elongator mutants is caused by a synergistic interaction between the *ssd1-d2* and *elp3Δ* alleles.

### The *ssd1-d2* allele augments phenotypes induced by the simultaneous lack of mcm^5^ and s^2^ groups

In the formation of mcm^5^s^2^U_34_, Elongator promotes synthesis of the mcm^5^ side-chain whereas the thiolation of the 2-position is catalyzed by the Ncs2/Ncs6 complex [40, 43, 53–55]. The simultaneous lack of mcm^5^ and s^2^ groups was originally reported to be lethal [40]. However, those experiments were performed in the W303 background and more recent studies have shown that strains lacking both groups are viable in the S288C background [18, 41]. To investigate if the allele at the *SSD1* locus accounts for the difference in viability, we constructed *ssd1-d2 elp3Δ ncs2Δ* and *SSD1 elp3Δ ncs2Δ* strains in both backgrounds all carrying a wild-type *ELP3* gene on a low-copy *URA3* plasmid. Analyses of the strains revealed that the *elp3Δ ncs2Δ* double mutant is viable in the W303 background if it encompasses a *SSD1* allele (Fig 4A). In the S288C background, the *ssd1-d2 elp3Δ ncs2Δ* strain is viable but it grows slower than the *SSD1 elp3Δ ncs2Δ* strain (Fig 4A; S5 Fig shows the same plates after 2 days of incubation). These observations not only show that allele at the *SSD1* locus influences the viability of *elp3Δ ncs2Δ* cells, but they also indicate that the growth phenotypes are modulated by additional genetic factors.

**Fig 4.**
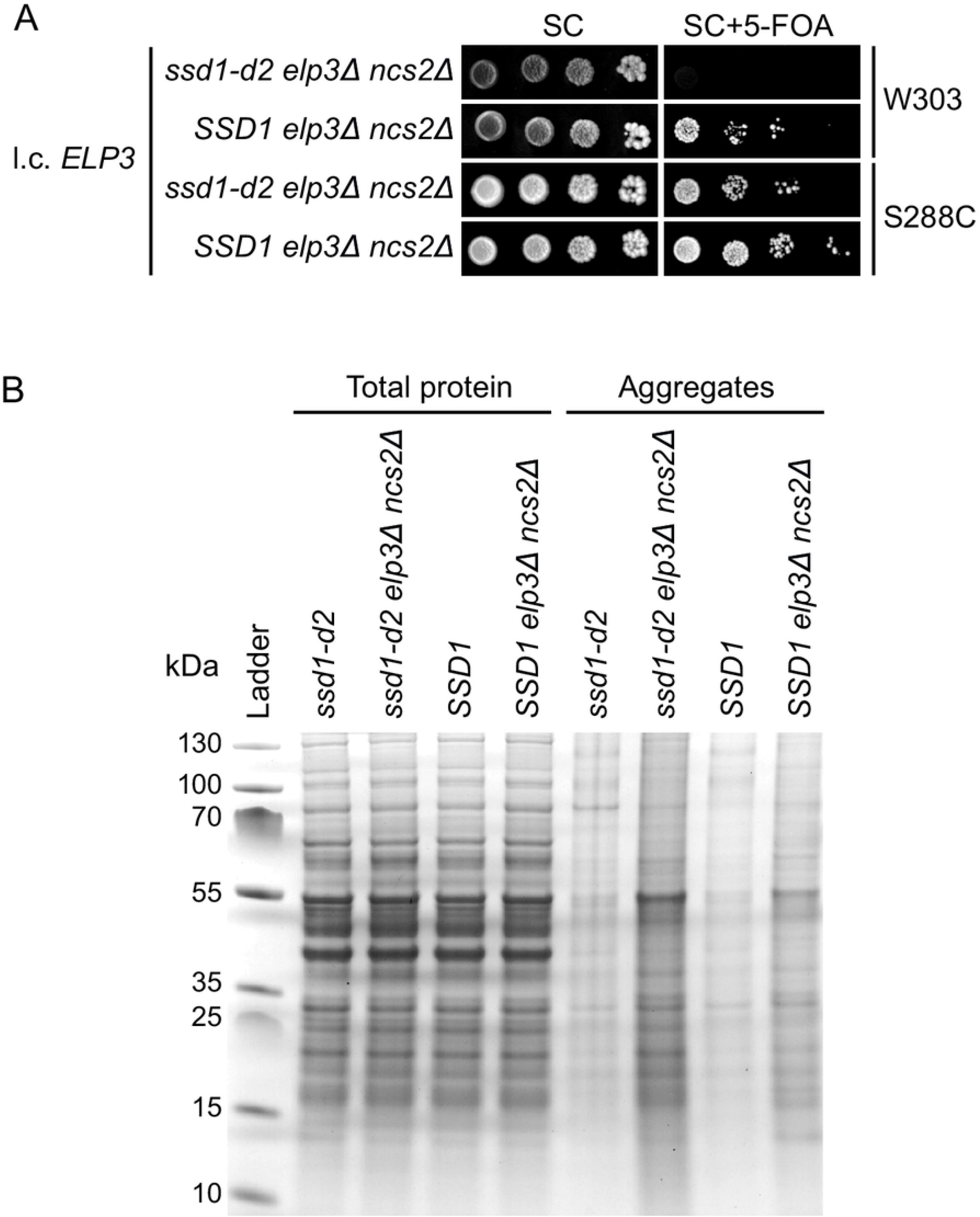
The growth and protein homeostasis defects of *elp3Δ ncs2Δ* cells are augmented by the *ssd1-d2* allele. (A) Influence of the *ssd1-d2* allele on the growth of *elp3Δ ncs2Δ* cells. The *ssd1-d2 elp3Δ ncs2Δ* (UMY4454 and MJY1159) and *SSD1 elp3Δ ncs2Δ* (UMY4467 and MJY1058) strains carrying the l.c. *URA3* plasmid pRS316-ELP3 were grown over-night at 30°C in SC medium, serially diluted, spotted on SC and SC+5-FOA plates, and incubated for 3 days at 30°C. (B) Effects of the *ssd1-d2* allele on protein aggregation in *elp3Δ ncs2Δ* cells. Total protein and protein aggregates was analyzed from the *ssd1-d2* (UMY4432), *ssd1-d2 elp3Δ ncs2Δ* (UMY4449), *SSD1* (BY4741) and *SSD1 elp3Δ ncs2Δ* (MJY1058) strains grown in SC medium at 30°C. The gel is a representative of two independent experiments.

The lack of the mcm^5^ and/or s^2^ groups has, in the S288C background, been shown to correlate with an increased accumulation of protein aggregates [18]. This effect was most pronounced in a strain lacking both groups and the phenotype was suggested to be a consequence of co-translational misfolding due to slower decoding of AAA and CAA codons by the hypomodified 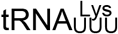 and 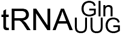 [18]. Moreover, the increased load of aggregates during normal growth was proposed to account for observation that the double mutant is impaired in clearing diamide-induced protein aggregates [18]. As strains deleted for *SSD1* show a defect in the disaggregation of heat-shock-induced protein aggregates [56], it seemed possible that the *ssd1-d2* allele would augment the protein homeostasis defect in *elp3Δ ncs2Δ* cells. To investigate this possibility, we isolated aggregates [18, 57] from the *ssd1-d2, SSD1, ssd1-d2 elp3Δ ncs2Δ*, and *SSD1 elp3Δ ncs2Δ* strains. Analyses of the insoluble fractions revealed that the levels of aggregated proteins are comparable in the *ssd1-d2* and *SSD1* strains (Fig 4B). However, the *ssd1-d2 elp3Δ ncs2Δ* mutant shows increased accumulation of aggregates compared to the *SSD1 elp3Δ ncs2Δ* strain (Fig 4B). Thus, the allele at the *SSD1* locus modulates the protein homeostasis defect induced by the simultaneous lack of the wobble mcm^5^ and s^2^ groups.

## Discussion

The phenotypic penetrance of a mutation is often impacted by the genetic background, a phenomenon frequently observed in monogenic diseases [58, 59]. In this study, we investigate the effect of genetic background on the phenotypes of *S. cerevisiae* mutants defective in the formation of modified wobble uridines in tRNA. We show that the phenotypes of Elongator mutants are augmented by the *ssd1-d2* allele found in some wild-type laboratory strains. Moreover, the histone H3 acetylation and telomeric gene silencing defects reported for Elongator mutants are only observed in cells harboring the *ssd1-d2* allele. Thus, the *ssd1-d2* allele sensitizes yeast cells to the effects induced by the lack mcm^5^/ncm^5^ groups in U_34_-containing tRNAs.

Although the pleiotropic phenotypes of Elongator mutants are largely caused by the reduced functionality of the hypomodified 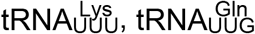 and 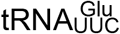, the basis for individual phenotypes is poorly understood. Several not necessarily mutually exclusive models have been proposed to explain how the lack of the mcm^5^/ncm^5^ groups can lead to a particular phenotype. One model postulates that phenotypes can be induced by inefficient translation of mRNAs enriched for AAA, CAA and/or GAA codons and the consequent effects on the abundance of the encoded factors [21, 60–62]. In this model, the inefficient decoding of the mRNAs leads to reduced protein output without affecting transcript abundance. The mechanism by which the slower decoding of the codons leads to reduced protein levels is unclear, but it may involve the inhibition of translation initiation by the queuing of ribosomes. Alternative models suggest that the phenotypes can be caused by defects in protein homeostasis and/or by indirect effects on transcription [18, 22]. The inactivation of *SSD1* not only influences translation and stability of the transcripts normally bound by Ssd1, but it also leads to altered abundance of many transcripts that do not appear to be Ssd1-associated [28, 39]. Thus, the effects of the *ssd1-d2* allele on the phenotypes of *elp3Δ* cells could be due to either direct or indirect effects on gene expression. Moreover, the ribosome profiling experiments of Elongator mutants have been performed in the S288C background [18, 22, 23] and it remains possible that the lack of Ssd1 influences the pausing at the AAA, CAA, and GAA codons.

While the difference at the *SSD1* locus partially explains the nonviability of *elp3Δ ncs2Δ* cells in the W303 background, the W303-derived *SSD1 elp3Δ ncs2Δ* strain grows slower than the corresponding strain in the S288C background (S5 Fig). Moreover, the *ssd1-d2 elp3Δ ncs2Δ* strain is viable, although with a growth defect, in the S288C background. These findings indicate that the growth phenotypes of *elp3Δ ncs2Δ* cells strains are modulated by additional genetic factors. Consistent with the finding that *ssd1Δ* cells show a defect in Hsp104-mediated protein disaggregation [56], the *ssd1-d2 elp3Δ ncs2Δ* strain shows increased accumulation of protein aggregates compared to the *SSD1 elp3Δ ncs2Δ* strain in the S288C background. It is, however, unclear if this increase is the cause or the consequence of the reduced growth of the *ssd1-d2 elp3Δ ncs2Δ* strain.

## Materials and Methods

### Yeast strains, plasmids, media and genetic procedures

Strains and plasmids used in this study are listed in S5 Table and S6 Table. Yeast media were prepared as described [63, 64]. The medium was where appropriate supplemented with 2.5 ng/ml rapamycin (R0395, Sigma-Aldrich), 7 mM caffeine (C0750, Sigma-Aldrich), 100 mM hydroxyurea (H8627, Sigma-Aldrich), 0.25 ng/ml diamide (D3648, Sigma-Aldrich), or 1 mg/ml 5-fluoroorotic acid (R0812, Thermo Fisher).

To generate *ssd1-d2* derivatives of BY4741 and BY4742 (S288C background)[65], we first replaced the sequence between position 2907 and 3315 of the *SSD1* ORF with a *URA3* gene PCR-amplified from pRS316 [66]. The oligonucleotides used for strain constructions are described in S7 Table. The generated strains were transformed with an *ssd1-d2* DNA fragment PCR-amplified from W303-1A [67]. Following selection on 5-fluoroorotic acid (5-FOA)-containing plates and subsequent single cell streaks, individual clones were screened for the integration of *ssd1-d2* allele by PCR and DNA sequencing. The generated strains (UMY4432 and UMY4433) were allowed to mate producing the homozygous *ssd1-d2/ssd1-d2* strain (UMY4434).

Strains deleted for *ELP3* or *NCS2* were constructed by transforming the appropriate diploid (UMY3387, UMY2836 or UMY4434) with an *elp3::KanMX4* or *ncs2::KanMX4* DNA fragment PCR-amplified from UMY3269 (*elp3::KanMX4*) or UMY3442 (*ncs2::KanMX4*) with appropriate homologies. Following PCR confirmation of the deletion, the generated heterozygous diploids were allowed to sporulate and the UMY4456 (*elp3Δ SSD1*, W303), MJY1019 (*ncs2Δ SSD1*, W303), UMY4439 (*elp3Δ ssd1-d2*, S288C), UMY4442 (*ncs2Δ ssd1-d2*, S288C), MJY1036 (*elp3Δ SSD1*, S288C), and MJY1021 (*ncs2Δ SSD1*, S288C) strains were obtained from tetrads. The *elp3Δ ncs2Δ SSD1* mutants (MJY1058 and UMY4467) were obtained from crosses between the relevant strains. The diploids used to generate the *elp3Δ ncs2Δ ssd1-d2* strains (MJY1159 and UMY4454) were transformed with pRS316-*ELP3* [40] before sporulation. MJY1159 was able to lose the plasmid generating strain UMY4449.

To construct plasmids carrying individual genes for factors in the CWI signaling pathway, we PCR-amplified the gene of interest using oligonucleotides that introduce appropriate restriction sites (S7 Table). The DNA fragment was then cloned into the corresponding sites of pRS425 [68] or pRS315 [66].

### RNA methods

The abundance of individual tRNA species was determined in total RNA isolated from exponentially growing cultures at an optical density at 600 nm (OD_600_) of ≈0.5 [64]. Samples containing 10 μg of total RNA were separated on 8M urea-containing 8% polyacrylamide gels followed by electroblotting to Zeta-probe membranes (Bio-Rad). The blots were sequentially probed for 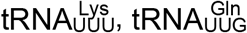 and 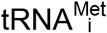 using ^32^P-labeled oligonucleotides (S7 Table). Signals were detected and analyzed by phosphorimaging using a Typhoon FLA 9500 biomolecular imager and Quantity One software.

To analyze the induction of *GAL1* transcripts, cells were grown at 30°C in 50 ml synthetic complete (SC) medium containing 2% raffinose (SC/Raf) to OD_600_≈0.45. The culture was harvested by centrifugation at 1,500 x g for 5 min at room temperature, and the cell pellet resuspended in 15 ml pre-warmed (30°C) SC/Raf medium. Following reincubation in the shaking water bath for 10 min, transcription of *GAL1* was induced by the addition of 1.5 ml pre-warmed 20% galactose. Aliquots were harvested [64] at various time points after the addition of galactose and the cell pellets frozen on dry ice. The procedures for determining mRNA levels have been described [64].

To analyze the nucleoside composition of total tRNA, the tRNA was isolated from exponentially growing cultures at OD_600_≈0.8 [21]. The tRNA was digested to nucleosides using nuclease P1 (Sigma-Aldrich, N8630) and bacterial alkaline phosphatase (Sigma-Aldrich, P4252) and the hydrolysate analyzed by HPLC [69, 70]. The compositions of the elution buffers were as described [70] with the difference that methanol concentration in buffer A was changed to 5% (v/v).

### Histone preparation and immunoblot analyses

Histones were isolated from cells grown in SC medium at 30°C to OD_600_≈0.8. Cells representing 100 OD_600_ units were harvested, washed once with water, resuspended in 30 ml of buffer A (0.1 mM Tris-HCl at pH 9.4, 10 mM DTT), and incubated on a rotator at 30°C for 15 min. Cells were collected, washed with 30 ml buffer B (1 M Sorbitol, 20 mM HEPES at pH7.4) and resuspended in 25 ml of buffer B containing 600 U yeast lytic enzyme. After 1 hour incubation on a rotator at 30°C, the sample was mixed with 25 ml of ice-cold buffer C (1 M Sorbitol, 20 mM PIPES at pH 6.8, 1 mM MgCl_2_) followed by centrifugation at 1,500 x g for 5 min. The pellet was resuspended in 40 ml nuclei isolation buffer [71] and the suspension incubated with gentle mixing at 4°C for 30 min. Cell debris were removed by centrifugation at 1,500 x g for 5 min. The supernatant was homogenized with 5 strokes in a Dounce homogenizer followed by centrifugation at 20,000 x g for 10 min. Histones in the pelleted nuclei were extracted by re-suspension in 5 ml of cold 0.2 M H_2_SO_4_ and overnight incubation on a rotator at 4°C. After centrifugation at 10,000 x g for 10 min, proteins in the supernatant were precipitated by adding 0.5 volumes of 100% trichloroacetic and 30 min of incubation on ice. Following centrifugation, the pellet was washed twice with acetone and then dissolved in 200 μl 10 mM Tris-HCl at pH 8.0. Fractions (10 μl) were resolved by 15% SDS-PAGE and transferred to Immobilon-P (Millipore) membranes. The blots were incubated with rabbit anti-acetyl-histone H3 (Lys14) antibodies (1:1,000 dilution, Millipore, 07-353) and then with horseradish peroxidase-linked donkey anti-rabbit IgG (NA934, GE Healthcare). Blots were stripped and reprobed with rabbit anti-histone H3 antibodes (1:5,000 dilution, Millipore, 07-690). Proteins were detected using ECL Western blotting detection reagents (GE Healthcare, RPN2209) and Amersham Hyperfilm ECL (GE Healthcare, 28906836).

### Analysis of protein aggregates

Protein aggregates were analyzed in exponentially growing cultures in SC medium at 30°C. Cells representing 50 OD_600_ units were harvested at OD_600_≈0.5 and protein aggregates were isolated [18, 57] from samples containing 5 mg of total protein. 1/10 of the aggregates and 5μg of total protein were resolved on a 4-12% NuPAGE Bis-Tris gel (Thermo Fisher, NP0321BOX) followed by staining with the Colloidal Blue Staining Kit (Thermo Fisher, LC6025).

## Acknowledgments

We thank members of M.J.’s and A.B.’s laboratories for valuable discussions.

## Supporting information

**S1 Fig.** The S288C-derived *elp3Δ* strain does not appear to have cell wall integrity defect. (A) Growth of the *elp3Δ* (MJY1036) strain carrying the indicated high-copy (h.c.) or low-copy (l.c.) *LEU2* plasmids. Cells were grown over-night at 30°C in liquid SC-leu medium, serially diluted, spotted on SC-leu plates, and incubated at 30°C or 37°C for 3 days. (B) The wild-type (W303-1A and BY4741) and *elp3Δ* (UMY3269 and MJY1036) strains were grown over-night at 30°C in liquid SC medium, serially diluted, spotted on SC plates and SC plates supplemented with 1M sorbitol. The plates were incubated for 3 days at 30°C or 37°C.

**S2 Fig.** HPLC analyses of the nucleoside composition of total tRNA from various *ssd1-d2* and *SSD1* strains. The peaks representing ncm^5^U, mcm^5^U, mcm^5^s^2^U, pseudouridine (ψ), cytidine (C), uridine (U), guanosine (G), and adenosine (A) are indicated. The asterisk indicates a peak that is a contamination from the bacterial alkaline phosphatase.

**S3 Fig.** Effects of *ssd1-d2* and *SSD1* alleles on the abundance of 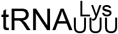 and 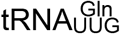. Northern analysis of total RNA isolated from the *ssd1-d2* (W303-1A and UMY4432), *ssd1-d2 elp3Δ* (UMY3269 and UMY4439), *SSD1* (UMY3385 and BY4741), and *SSD1 elp3Δ* (UMY4456 and MJY1036) strains grown in SC medium at 30°C. The blot was probed for 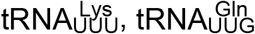, and 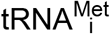 using radiolabeled oligonucleotides.

**S4 Fig.** The transcriptional activation of *GAL1* is not impaired in *elp3Δ* mutants. Northern analysis of total RNA isolated from the *ssd1-d2* (W303-1A), *ssd1-d2 elp3Δ* (UMY3269), *SSD1* (UMY3385) and *SSD1 elp3Δ* (UMY4456) strains. Cells were grown in SC medium containing 2% raffinose followed by induction of *GAL1* transcription by the addition of 0.1 volumes 20% galactose. Time points after the addition of galactose are indicated above the lanes. The blot was probed for *GAL1* transcripts using a randomly labelled DNA fragment. 18S rRNA was detected using a oligonucleotide probe. The blot is a representative of two independent experiments.

**S5 Fig.** Influence of the *ssd1-d2* allele on the growth of *elp3Δ ncs2Δ* cells. The figure shows a shorter incubation (2 days) of the plates in Fig. 4A.

**S1 Table.** Steady-state tRNA levels in *elp3Δ* cells carrying the indicated plasmids.

**S2 Table.** Relative amounts of ncm^5^U, mcm^5^U and mcm^5^s^2^U in total tRNA isolated from various strains.

**S3 Table.** Generation times of indicated strains grown at 30°C or 37°C.

**S4 Table.** Steady-state tRNA levels in the indicated strains.

**S5 Table.** Yeast strains used in this study.

**S6 Table.** Plasmids used in this study.

**S7 Table.** Oligonucleotides used in this study.

