## Supplemental Data for "*SSD1* suppresses phenotypes induced by the lack of Elongator-dependent tRNA modifications"

S1 Fig

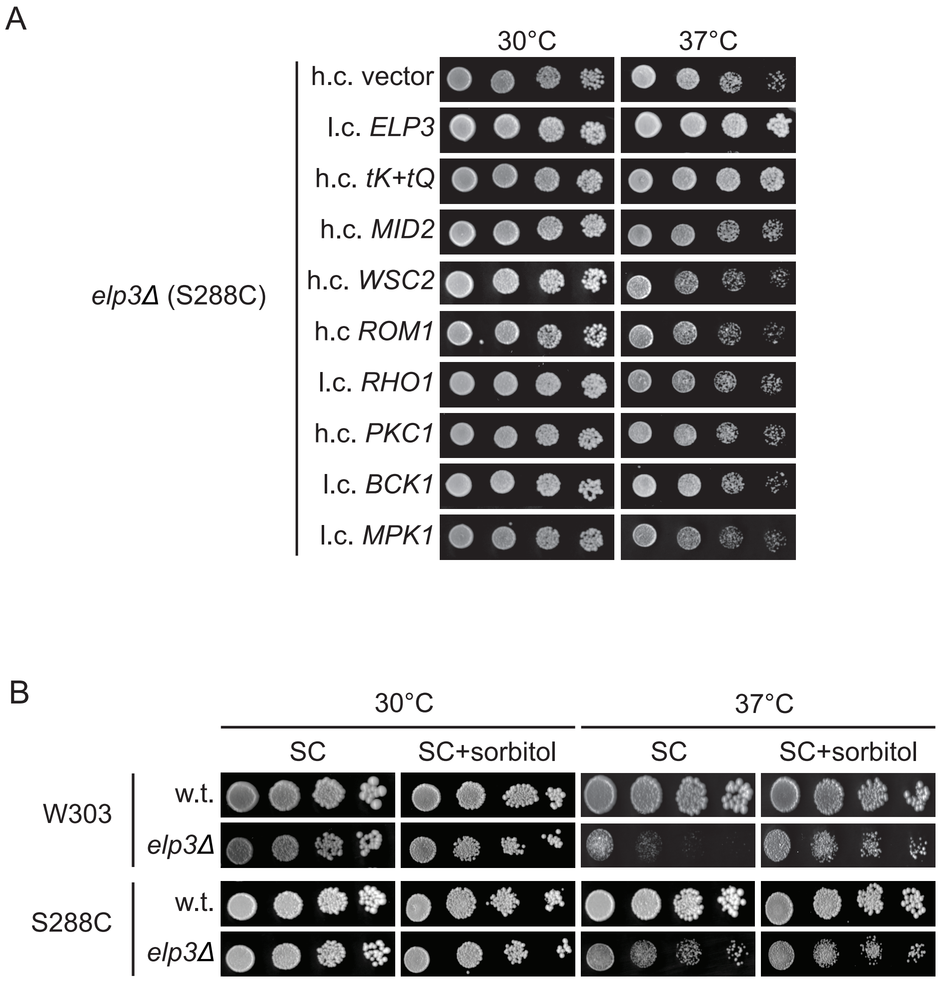
A
30°C
37°C
h.c. vector
l.c. ELP3
h.c. tK+tQ
h.c. MID2
h.c. WSC2
h.c ROM1
l.c. RHO1
h.c. PKC1
l.c. BCK1
l.c. MPK1
elp3Δ (S288C)
B
30°C
37°C
SC
SC+sorbitol
SC
SC+sorbitol
W303
w.t.
elp3Δ
S288C
w.t.
elp3Δ

S2 Fig

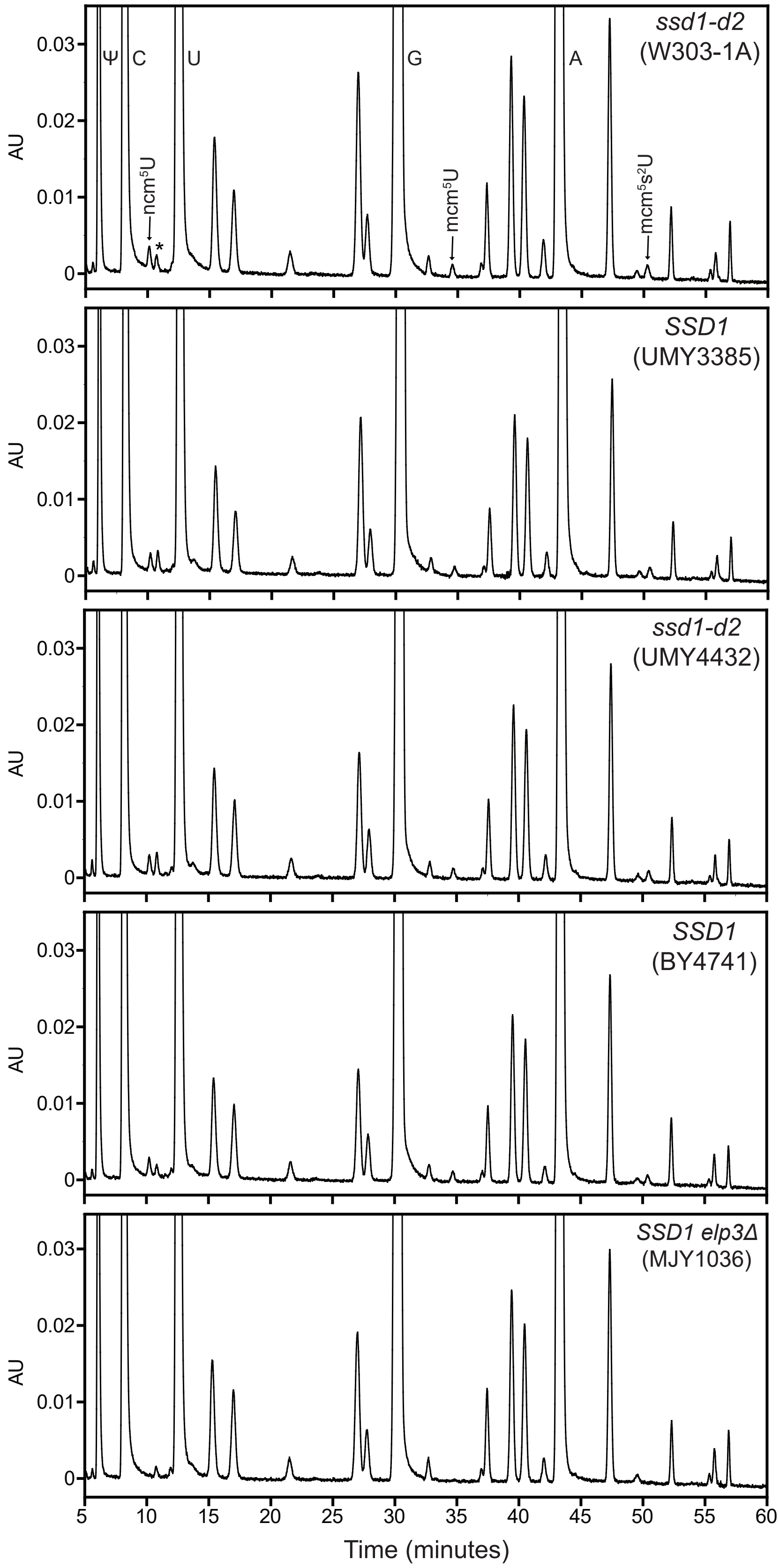

ssd1-d2
(W303-1A)
Ψ
C
U
G
A
ncm5U
*
mcm5U
mcm5s2U
SSD1
(UMY3385)
ssd1-d2
(UMY4432)
SSD1
(BY4741)
SSD1 elp3Δ
(MJY1036)
AU
0.03
0.02
0.01
0
5
10
15
20
25
30
35
40
45
50
55
60
Time (minutes)

S3 Fig

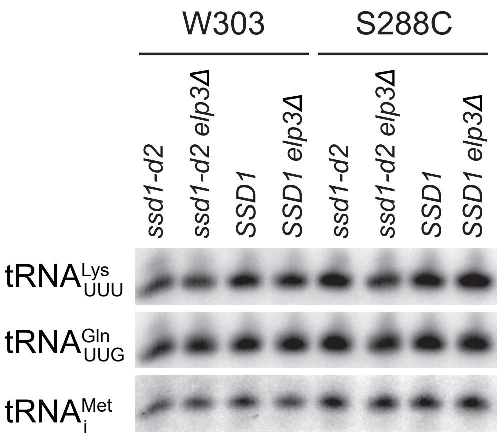
W303
S288C
ssd1-d2
ssd1-d2 elp3Δ
SSD1
SSD1 elp3Δ
ssd1-d2
ssd1-d2 elp3Δ
SSD1
SSD1 elp3Δ
tRNA Lys UUU
tRNA Gln UUG
tRNA Met i

S4 Fig

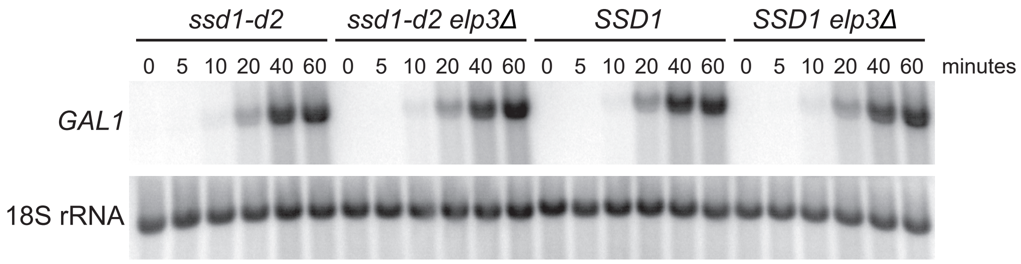

ssd1-d2
ssd1-d2 elp3Δ
SSD1
SSD1 elp3Δ
0 5 10 20 40 60 0 5 10 20 40 60 0 5 10 20 40 60 0 5 10 20 40 60 minutes
GAL1
18S rRNA

S5 Fig

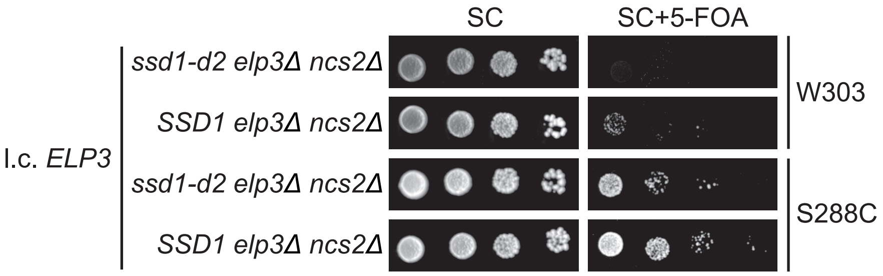

SC
SC+5-FOA
l.c. *ELP3*
*ssd1-d2 elp3Δ ncs2Δ*
*SSD1 elp3Δ ncs2Δ*
*ssd1-d2 elp3Δ ncs2Δ*
*SSD1 elp3Δ ncs2Δ*
W303
S288C

**S1 Table.** Steady-state tRNA levels in *elp3Δ* cells carrying the indicated plasmids.

| Plasmid | Relative tRNA level[a] | | | |
|---|---|---|---|---|
| | $tRNA^{Lys}_{UUU}$ | | $tRNA^{Gln}_{UUG}$ | |
| | 30°C | 37°C | 30°C | 37°C |
| h.c. vector | 1.00 | 1.00 | 1.00 | 1.00 |
| l.c. *ELP3* | 1.44 ± 0.07 | 1.44 ± 0.45 | 1.03 ± 0.05 | 1.02 ± 0.12 |
| h.c. *tK(UUU)-tQ(UUG)* | 2.12 ± 0.39 | 2.49 ± 0.67 | 2.19 ± 0.39 | 2.84 ± 0.22 |
| h.c. *PKC1* | 0.97 ± 0.22 | 1.22 ± 0.36 | 0.98 ± 0.19 | 1.06 ± 0.19 |

[a] The signal for the indicated tRNA species was normalized to the corresponding $tRNA^{Met}_{i}$ signal and the value expressed relative to that for the strain carrying the empty h.c. vector. The values represent the average from the blot shown in Fig 1B and two additional independent experiments. The standard deviation is indicated.

**S2 Table.** Relative amounts of ncm$^5$U, mcm$^5$U and mcm$^5$s$^2$U in total tRNA isolated from various strains.

| Background | Strain | Modified nucleoside[a] | | |
|---|---|---|---|---|
| | | ncm$^5$U | mcm$^5$U | mcm$^5$s$^2$U |
| W303 | *ssd1-d2* (W303-1A) | 1.00 | 1.00 | 1.00 |
| | *SSD1* (UMY3385) | 0.95 ± 0.08 | 1.00 ± 0.21 | 0.87 ± 0.13 |
| | *ssd1-d2 elp3Δ* (UMY3269) | -[b] | - | - |
| | *SSD1 elp3Δ* (UMY4456) | - | - | - |
| S288C | *ssd1-d2* (UMY4432) | 0.99 ± 0.07 | 1.18 ± 0.16 | 0.86 ± 0.11 |
| | *SSD1* (BY4741) | 0.97 ± 0.08 | 1.01 ± 0.22 | 0.83 ± 0.17 |
| | *ssd1-d2 elp3Δ* (UMY4439) | - | - | - |
| | *SSD1 elp3Δ* (MJY1036) | - | - | - |

[a] The peaks for ncm$^5$U, mcm$^5$U and mcm$^5$s$^2$U were integrated and the values normalized to the value for pseudouridine (Ψ). The normalized values were expressed relative to the corresponding values from the W303-1A strain, which was set to 1. The values represent the average of three independent experiments and their standard deviations.

[b] Below the detection limit.

**S3 Table.** Generation times of indicated strains grown at 30°C or 37°C.

| Background | Strain | Generation time (h)[a] | |
|---|---|---|---|
| | | 30°C | 37°C |
| W303 | *ssd1-d2* (W303-1A) | 1.67 ± 0.10 | 1.96 ± 0.13 |
| | *SSD1* (UMY3385) | 1.61 ± 0.12 | 1.84 ± 0.06 |
| | *ssd1-d2 elp3Δ* (UMY3269) | 2.32 ± 0.20 | 3.53 ± 0.21 |
| | *SSD1 elp3Δ* (UMY4456) | 1.99 ± 0.07 | 2.69 ± 0.21 |
| S288C | *ssd1-d2* (UMY4432) | 1.47 ± 0.02 | 1.91 ± 0.16 |
| | *SSD1* (BY4741) | 1.43 ± 0.03 | 1.61 ± 0.04 |
| | *ssd1-d2 elp3Δ* (UMY4439) | 2.26 ± 0.11 | 3.66 ± 0.22 |
| | *SSD1 elp3Δ* (MJY1036) | 2.11 ± 0.04 | 2.68 ± 0.12 |

[a] Growth rates were determined in SC medium. The values represent the average from four independent experiments and their standard deviations.

**S4 Table.** Steady-state tRNA levels in the indicated strains.

| Background | Strain | Relative tRNA level[a] | |
|---|---|---|---|
| | | $tRNA^{Lys}_{UUU}$ | $tRNA^{Gln}_{UUG}$ |
| W303 | *ssd1-d2* (W303-1A) | 1.00 | 1.00 |
| | *ssd1-d2 elp3Δ* (UMY3269) | 0.93 ± 0.04 | 1.09 ± 0.24 |
| | *SSD1* (UMY3385) | 1.24 ± 0.17 | 1.15 ± 0.16 |
| | *SSD1 elp3Δ* (UMY4456) | 1.14 ± 0.06 | 1.19 ± 0.23 |
| S288C | *ssd1-d2* (UMY4432) | 1.00 | 1.00 |
| | *ssd1-d2 elp3Δ* (UMY4439) | 0.84 ± 0.06 | 1.01 ± 0.09 |
| | *SSD1* (BY4741) | 1.03 ± 0.07 | 1.03 ± 0.07 |
| | *SSD1 elp3Δ* (MJY1036) | 1.09 ± 0.09 | 1.03 ± 0.14 |

[a] The signal for the indicated tRNA species was normalized to the corresponding $tRNA^{Met}_{i}$ signal and the value expressed relative to that for the respective *ssd1-d2* strain, which was set to 1. The values represent the average from the blot shown in S3 Fig and two additional independent experiments. The standard deviation is indicated.

**S5 Table.** Yeast strains used in this study.

| Strain | Genotype | Source or reference |
|---|---|---|
| BY4741 | *MATa his3Δ1 leu2Δ0 met15Δ0 ura3Δ0 SSD1* | [65] |
| BY4742 | *MATα his3Δ1 leu2Δ0 lys2Δ0 ura3Δ0 SSD1* | [65] |
| UMY2836 | Diploid between BY4741 and BY4742 | This lab |
| MJY1021 | *MATα his3Δ1 leu2Δ0 lys2Δ0 ura3Δ0 SSD1 ncs2::kanMX4* | This study |
| MJY1036 | *MATa his3Δ1 leu2Δ0 met15Δ0 ura3Δ0 SSD1 elp3::kanMX4* | This study |
| MJY1058 | *MATa his3Δ1 leu2Δ0 met15Δ0 ura3Δ0 SSD1 elp3::kanMX4 ncs2::kanMX4* | This study |
| MJY1159 | *MATa his3Δ1 leu2Δ0 met15Δ0 ura3Δ0 ssd1-d2 elp3::KanMX4 ncs2::kanMX4* pRS316-*ELP3* | This study |
| UMY4449 | *MATa his3Δ1 leu2Δ0 met15Δ0 ura3Δ0 ssd1-d2 elp3::kanMX4 ncs2::kanMX4* | This study |
| UMY4432 | *MATa his3Δ1 leu2Δ0 met15Δ0 ura3Δ0 ssd1-d2* | This study |
| UMY4433 | *MATα his3Δ1 leu2Δ0 lys2Δ0 ura3Δ0 ssd1-d2* | This study |
| UMY4434 | Diploid between UMY4432 and UMY4433 | This study |
| UMY4439 | *MATa his3Δ1 leu2Δ0 met15Δ0 ura3Δ0 ssd1-d2 elp3::kanMX4* | This study |
| UMY4442 | *MATα his3Δ1 leu2Δ0 lys2Δ0 ura3Δ0 ssd1-d2 ncs2::kanMX4* | This study |
| UMY4445 | Diploid between UMY4439 and UMY4442 | This study |
| W303-1A | *MATa leu2-3,112 trp1-1 can1-100 ura3-1 ade2-1 his3-11,15 ssd1-d2* | [67] |
| W303-1B | *MATα leu2-3,112 trp1-1 can1-100 ura3-1 ade2-1 his3-11,15 ssd1-d2* | [67] |
| UMY3269 | *MATa leu2-3,112 trp1-1 can1-100 ura3-1 ade2-1 his3-11,15 ssd1-d2 elp3::kanMX4* | [21] |
| UMY3442 | *MATα leu2-3,112 trp1-1 can1-100 ura3-1 ade2-1 his3-11,15 ssd1-d2 ncs2::kanMX4* | [20] |
| UMY4444 | Diploid between UMY3269 and UMY3442 | This study |
| UMY3385 | *MATa leu2-3,112 trp1-1 can1-100 ura3-1 ade2-1 his3-11,15 SSD1* | [43] |
| UMY3386 | *MATα leu2-3,112 trp1-1 can1-100 ura3-1 ade2-1 his3-11,15 SSD1* | [43] |
| UMY3387 | Diploid between UMY3385 and UMY3386 | [43] |
| MJY1019 | *MATα leu2-3,112 trp1-1 can1-100 ura3-1 ade2-1 his3-11,15 SSD1 ncs2::kanMX4* | This study |
| UMY4456 | *MATa leu2-3,112 trp1-1 can1-100 ura3-1 ade2-1 his3-11,15 SSD1 elp3::kanMX4* | This study |
| UMY4467 | *MATa leu2-3,112 trp1-1 can1-100 ura3-1 ade2-1 his3-11,15 SSD1 elp3::kanMX4 ncs2::kanMX4* | This study |
| UMY4454 | *MATa leu2-3,112 trp1-1 can1-100 ura3-1 ade2-1 his3-11,15 ssd1-d2 elp3::kanMX4 ncs2::kanMX4 pRS316-ELP3* | This study |
| UMY2584 | *MATa leu2-3,112 trp1-1 can1-100 ura3-1 ade2-1 his3-11,15 ssd1-d2 TELVIIL::URA3 TELVR::ADE2* | [21] |
| UMY3790 | *MATa leu2-3,112 trp1-1 can1-100 ura3-1 ade2-1 his3-11,15 ssd1-d2 TELVIIL::URA3 TELVR::ADE2 elp3::kanMX4* | [21] |

**S6 Table.** Plasmids used in this study

| Plasmid | Description | Source or reference |
|---|---|---|
| pRS316 | YCp, *URA3* | [66] |
| pRS315 | YCp, *LEU2* | [66] |
| pRS425 | YEp, *LEU2* | [68] |
| pABY2160 | pRS425-*MID2* | This study |
| pMJ120 | pRS425-*WSC2* | This study |
| pABY2176 | pRS425-*ROM1* | This study |
| pABY2128 | pRS425-*RHO1* | This study |
| pABY2177 | pRS315-*RHO1* | This study |
| pABY2129 | pRS425-*PKC1* | This study |
| pABY2130 | pRS425-*BCK1* | This study |
| pABY2239 | pRS315-*BCK1* | This study |
| pABY2131 | pRS425-*MPK1* | This study |
| pABY2240 | pRS315-*MPK1* | This study |
| pABY2227 | pRS315-*SSD1* | This study |
| pABY1707 | pRS425-*tK(UUU)-tQ(UUG)* | [72] |
| pABY1514 | pRS315-*ELP3* | [5] |
| pABY1481 | pRS316-*ELP3* | [40] |

**S7 Table.** Oligonucleotides used in this study

| Name | Purpose | Sequence | Restriction site |
|---|---|---|---|
| o3258 | *MID2* cloning | AAAAACTAGTTTGCTTTCATAATCTGCAAAT | *Spe*I |
| o3259 | *MID2* cloning | AAAACCCGGGCAGTGGAACGTTAAAGCACT | *Sma*I |
| o2945 | *WSC2* cloning | AAAAGCGGCCGCTTGTGATCTAGCACTTCTC | *Not*I |
| o2946 | *WSC2* cloning | AAAAGTCGACGTAATGTGGAGATCATCG | *Sal*I |
| o3248 | *ROM1* cloning | AAAAACTAGTACTTTTGCCATCTTATACTCATC | *Spe*I |
| o3249 | *ROM1* cloning | AAAACCCGGGCTTCAATACGGTCAGAATTATC | *Sma*I |
| o3106 | *RHO1* cloning | AAAAGCGGCCGCAGGTTGGGTTATGGAACCTT | *Not*I |
| o3107 | *RHO1* cloning | AAAAGTCGACTTGCCAGGTGTTAAGAAGG | *Sal*I |
| o3104 | *PKC1* cloning | AAAAGCGGCCGCCGCGAACTCGTAAGTAGAAAA | *Not*I |
| o3105 | *PKC1* cloning | AAAAGTCGACACATTCTTCAAATGCCTGC | *Sal*I |
| o3118 | *BCK1* cloning | AAAACCCGGGTCCATATTTGGTGACCGA | *Sma*I |
| o3119 | *BCK1* cloning | AAAAGAGCTCGAGGACCTTCCTGATGAAAG | *Sac*I |
| o3102 | *MPK1* cloning | AAAAGCGGCCGCGAGCGGTAACTATGGACACC | *Not*I |
| o3103 | *MPK1* cloning | AAAAGTCGACGAGTACGATTAAGATAAGCGTCG | *Sal*I |
| o2975 | *SSD1* cloning | AAAAGGATCCTCACGAGTATTTTCGCTC | *Bam*HI |
| o2976 | *SSD1* cloning | AAAAGAGCTCCGGAAAAATTACCCAGC | *Sac*I |
| o797 | *elp3::kanMX4* amplification | GCTTACACTTCGTTCCTTCC | |
| o798 | *elp3::kanMX4* amplification | CAGTGAGAGAAGGAGAAAGC | |
| o800 | *elp3Δ* confirmation | CATGTACGGTCGCTTGAGGT | |
| o799 | *elp3Δ* confirmation | CGTGCAATTGACCGAACGTG | |
| o3472 | *ssd1::URA3* amplification | CGTTGGCCAATCACATCTTTGCATCCATTTGGTA TTTTAGTGATGACGGTGAAAACCTCT | |
| o3473 | *ssd1::URA3* amplification | AACCGACAGCGTGGCTGATTCCTTGCCAGGGGC CAACGACCGGCCTATTGGTTAAAAAATG | |
| o3469 | *ssd1-d2* amplification | ATGTCGTTGCTGTTTTGGAC | |
| o3478 | *ssd1-d2* amplification | CCTAAAGTTCTATCCAGGATG | |
| o1423 | *ncs2::kanMX4* amplification | GATCTTTTCCACTGGTCGTC | |
| o1424 | *ncs2::kanMX4* amplification | CTACTTAAAGCCCAAGCCTC | |
| o1991 | *ncs2Δ* confirmation | TGGTATCGGTCTGCGATTCC | |
| o1425 | *ncs2Δ* confirmation | CTACGTCGACGGTGAGTGGTGGAGTTCCTC | |

| | | |
|---|---|---|
| o1128 | $tRNA^{Lys}_{UUU}$ probe | CCCTGACATTTCGGTTAA |
| o1439 | $tRNA^{Gln}_{UUG}$ probe | CCACTACACTATAGGACC |
| oMJ561 | $tRNA^{Met}_{i}$ probe | GGACATCAGGGTTATGAGCC |
